# Development of a high-throughput radial migration device

**DOI:** 10.1101/2020.01.31.928879

**Authors:** C. Ryan Oliver, Andrew C. Little, Trisha M. Westerhof, Pragathi Pathanjeli, Joel A. Yates, Sofia D. Merajver

**Affiliations:** Department of Internal Medicine, University of Michigan, Ann Arbor, MI, 48109, USA; Department of Biomedical Engineering, University of Michigan, Ann Arbor, MI, 48109, USA

## Abstract

By combining the radial migration assay with injection molded gaskets and a rigid fixture, we have developed a more reliable and sensitive method for measuring radial cell migration. This method is well adapted for use on high throughput automated imaging systems. The use of injection molded gaskets enables low cost replacement of cell-wetted components. Furthermore, the design enables secondary placement of attractants and co-cultures. This device and high-throughput application permit the use of therapeutic screening to evaluate phenotypic responses e.g. cancer cell migration. This approach is orthogonal to other 2D cell migration applications such as scratch wound assays, although here we offer a non-invasive, high-throughput device which is currently not commercially available. Collectively, we have designed a systematic, reliable, high-throughput application to monitor phenotypic responses to chemotherapeutic screens, genetic alterations (e.g. RNAi; CRISPR; others), supplemental regiments, and other approaches offering a reliable methodology to survey unbiased and non-invasive cell migration.

## INTRODUCTION

Many biological questions are concerned with the rate(s) of cellular migration, e.g. tissue development, cancer metastasis, immune cell trafficking, among many others. Understanding the precise mechanisms by which secreted factors from cellular microenvironments and the proteins which regulate downstream migratory responses are critical aspects of cellular biology (1). While many assays exist, one valuable assay is the radial migration assay (2); first introduced by Gilles et al., in 1999 (3). It has been developed into a considerable number of variations complete with novel methodologies and data processing techniques (4), including extracellular matrix deposition for 3D invasion assays among others (5–7). However, adoption of this technique is relatively slow because placement of the gasket for high throughput studies is tedious. Therefore, it is prudent to adapt the radial migration assay for high throughput applications by reducing the complexity and time necessary to setup and perform the assay.

One of the most popular 2D assays is the scratch wound/wound closure assay (8). This assay was introduced by Todaro et al. the late 1960s (9) and has been adapted and applied in many variations since (10). This assay has gained popularity due to advances in the tools available that create precise linear “wounds” in a 24-well plate using a single mechanical lever (11). One of the disadvantages to this technique is that creating a linear scratch (wound) in a cell monolayer initiates the release of inflammatory cytokines/chemokines and wound signals, undoubtedly influencing cellular responses and triggering cellular communication programs. Indeed, this can confound results in fields such as cancer where researchers desire an understanding of intrinsic migratory capacity free of influence from wound healing or damage responses (i.e. damage-associated molecular patterns (DAMPs; alarmins, others, see (12, 13))). Therefore, the radial migration assay offers an attractive alternative which permits the evaluation of 2D multidirectional, undirected cell migration, yet permits communication of bulk cells to leading edge migratory cells; modeling tumor dynamics in a 2D fashion.

Independent of migration assays, robust imaging techniques have been developed for tracking the migration of cells from precise location(s) (14). These techniques acquire an initial image that represents the diameter of the starting population and tracks the shape and growth in diameter of the cells as they migrate radially from a central point. Permitting the measurement of either bulk cellular migration or single cell dynamics. This data allows researchers to probe the relationship between phenotypic behavior and cellular pathways. Coupling high-throughput migration assays with automated imaging provides an attractive avenue to evaluate intrinsic cell mobility and further, evaluate the potential use of targeted therapies to slow or inhibit cell migration reproducibly. A highly reproducible assay allows for fewer experiments to be performed to arrive at convincing and robust conclusions, saving time and resources. This is particularly useful in the design of high value chemotherapeutics that aim to inhibit cancer metastasis or understand the relationship between genetic mediators of cancer cell migration or invasion; examples can be seen here (15, 16).

Here, we describe our newly designed radial migration device which enables rapid and precise application of the radial migration assay for fundamental and translational biology in absence of inducing a damage or inflammatory response. Our high-throughput radial migration tool (Fig. 1A) is a remarkably simple fixture utilizing sound engineered design principles. This tool allows for the seeding of cells at 2.6 mm diameters with 0.1 mm precision. Initially, we describe the method and device design, then provide modeling of the forces and characterization of the approach. Finally, we demonstrate its use to study the migration of cancer cells as a proof of principle. Collectively, we believe this new tool permits high-throughput adaptation of a previously tedious and time-consuming assay. When paired with high content imaging (e.g. BioTek BioSpa/Cytation5 incubator/imaging platform or IncuCyte Live-Cell Analysis platforms), this new assay will provide valuable information regarding unabated cell migration and offers a novel method to evaluate migratory responses.

**Figure 1 –.**
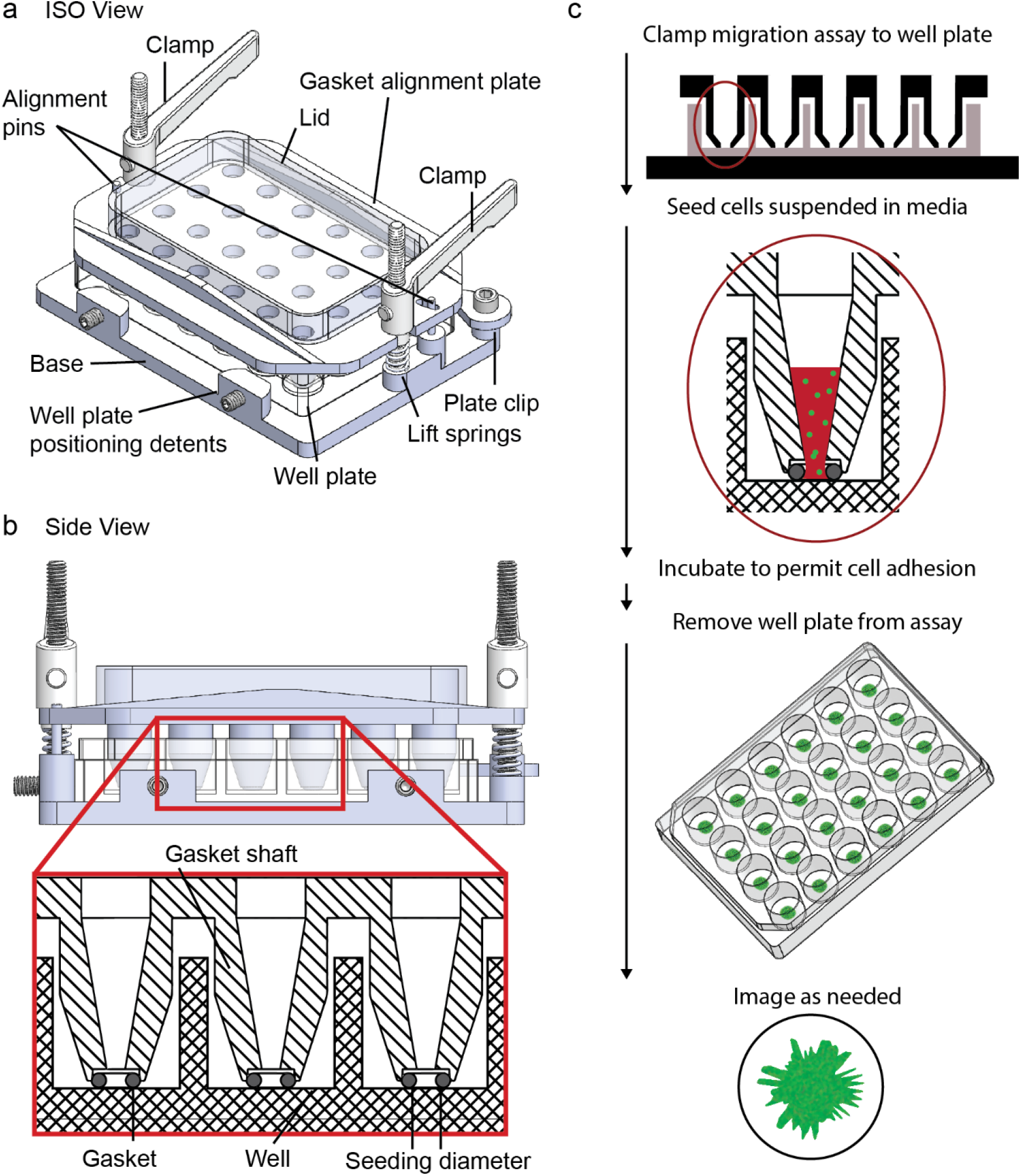
Radial migration assay overview and model application. a) ISO standard view of the radial migration assay as assembled. b) Side view of the radial migration device as assembled. Inset in red shows the placement of the gaskets between the well plate and the fixture. c) Standard protocol for using the radial migration assay to seed cells into a well plate.

## RESULTS

### Device Implementation and Experimental Approach

The approach we present here (Fig. 1A, Fig. 1B) is simply designed to alleviate barriers in the use and interpretation of radial migration assays. The method uses an aluminum fixture made of two parts—a base and a gasket alignment plate. The gasket alignment plate, which harbors an extruded shaft, aligns using steel pins and applies pressure to the gasket within each well of a 24 well plate. Commercially available gaskets (005 Silicone, McMaster-Carr) are suitable for this application and are inserted into recesses in the bottom of the locating shafts. Alternatively, custom gaskets can be created for more advanced studies. Clamps are located on both ends of the device and apply pressure between the upper and lower plate. Alignment detents are used for three-point kinematic locating of the well plate on the base plate.

To use the device, a 24-well cell culture plate is placed against the detents (Fig. 1C). Then the gasket alignment plate is clamped against the well plate. For sterility, the entire system, including the gaskets, can be autoclaved, cleaned with ethanol, or plasma cleaned. After being assembled, the central opening of each well is filled with 100 μL of cell culture media and placed in a vacuum Bell Jar with applied vacuum (10-100 mTorr) for ~2-3 min to remove air bubbles at the bottom of each well. Following vacuum, the device can now be loaded with cells to the existing media in each well. While cell concentration should be empirically determined, we find that ~50k cells per well is sufficient to form a confluent monolayer of cells in each well (Fig. 3F). After loading cells, a lid covers the openings for sterility and humidity control, and the device can be placed in the incubator to allow cells to become adherent. After the cells have adhered, the clamps can be released, and the gasket alignment plate can be removed. Springs will lift the gaskets and gasket alignment plate off the well plate in a uniform way to minimize peeling and other damage to the radial cell monolayer. Non-adherent cells and debris should be removed by 3-5 successive PBS washes. At this time cells may be supplemented with media components, drug treatments, or the cell monolayer may be deposited with Matrigel to observe 3D matrix invasion. Following cell treatment, the user should then initiate imaging to observe migratory/invasive responses as a function of time.

### Modeling Gasket Sealing Forces

To ensure proper and uniform sealing of the gaskets for each well, we combined a finite element analysis (FEA) model with accepted practices used in face sealing gaskets. First, considering the model for sealing a single gasket, this type of seal is an axial static face seal with a target of 20% compression. For a round gasket with a cross sectional diameter of 1.78 mm, and a circumference of 13.6 mm the force per linear mm applied to the seal can be tabulated from a standard table. For a shore hardness of 50A, the expected range is between 0.79 N/mm and 2.45 N/mm. We chose the value of 2.45 N/mm for a shore hardness of 50A because the compressive moduli of silicone is higher than other elastomeric materials. This results in a force of or 33.4 N per gasket. Because we are sealing 24 wells, the total force for the plate is approximately 802.8 N (7.5 MPa). Therefore, the applied load from each of the two clamps must be 401.4 N, and the selected ¼-20 bolts are rated for a clamping force of 5870 N. The fixtures were CNC machined and lightly bead blasted resulting in a 0.8 μm finish. These estimated forces should seal against the surface roughness of 0.8 μm specified on the gasket alignment plate. Finally, the inner gland/inset that houses the gaskets uses a 5% interference fit between the gland and the gasket. Given the thickness of the gasket at 1.78 mm and a compression of 0.38 mm, the displacement represents an additional 3%. This would require a nominal amount of additional pressure and lead towards a balance of deflection and compression.

**Figure 1 –.**
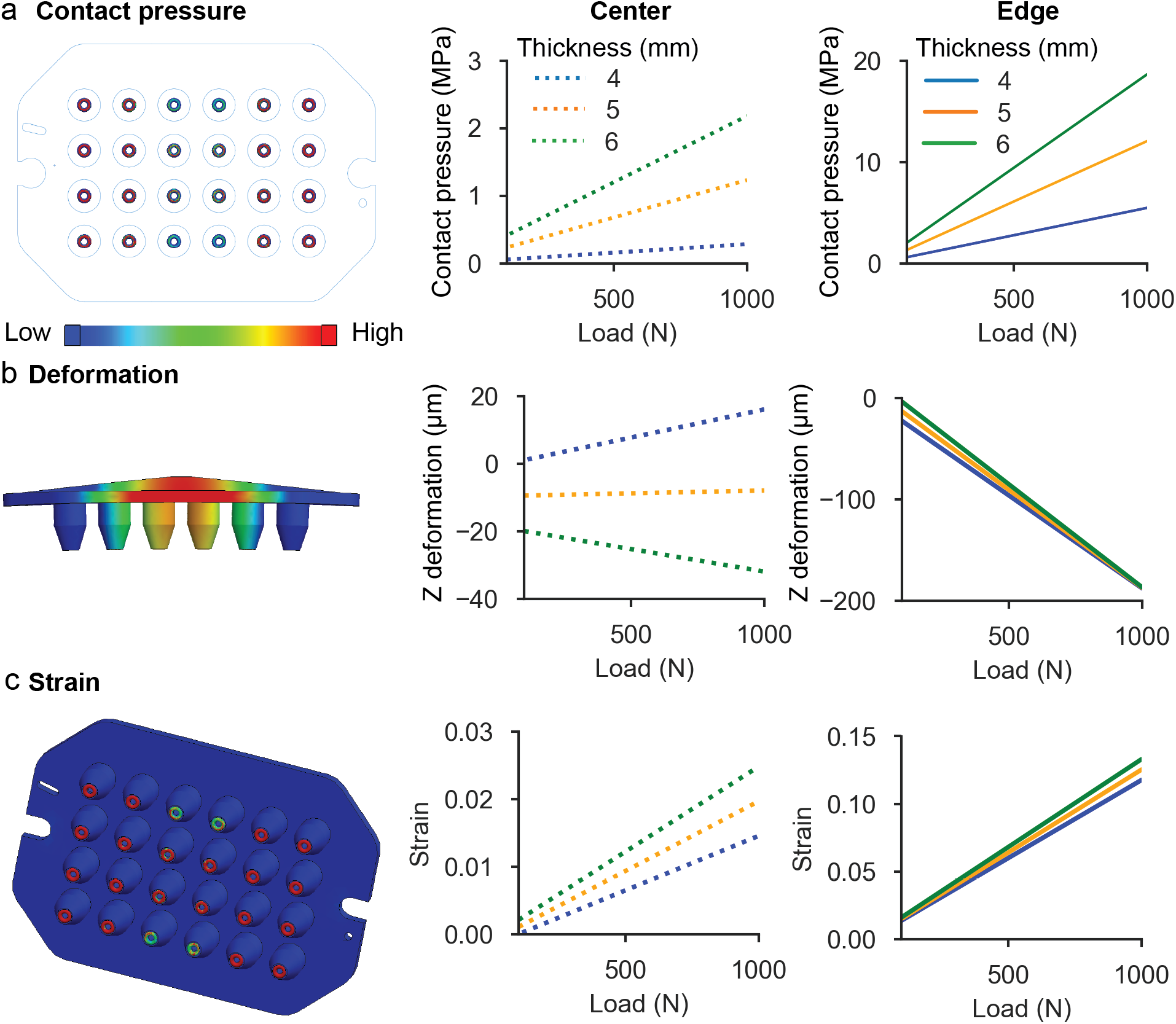
FEA simulation of forces and displacements under load. a) Distribution of contact pressure across the surface of the 24 well plate for 100 N load, 5 mm thickness (left). Red indicates higher pressures and blue lower. Plot of the contact pressure under varying loads in the center (middle) and edge (right) cases. b) Deformation profile of the fixture under a load of 100 N, 5 mm thickness (left). Red indicates larger deformations and blue lower. Plot of the gasket deformation under varying loads in the center (middle) and edge (right) cases. c) Strain distribution across the surface of the 24 well plate for a load of 100 N, 5 mm thickness (left). Red indicates higher strains and blue lower. Plot of the gasket strain under varying loads in the center (middle) and edge (right) cases.

To then understand how the force transferred from the gasket alignment plate varies between the gaskets and the well plate, an FEA model was used to simulate a range of loads and fixture thicknesses. The FEA model was performed using Solidworks built-in FEA system. The applied analysis was determined for a static force applied by the two clamps. The solver was a direct sparse matrix solver, h-adaptive. A non-penetrating contact constraint was applied between the gaskets, and the top and bottom fixture, as well as between both fixtures. Aluminum 6061 was specified as the material for the fixtures and a 50A shore durometer hardness variant of silicone rubber was specified for the gaskets to match the gaskets purchased for experimentation.

Different loads were applied to the model to observe the impact on the sealing/contact pressure between the gaskets, fixtures, and their deformation. In addition, the model was used to understand how the effective thickness of fixture can be optimized to reduce the bending curvature in the device due to the end loads which would prevent sealing in the center of the device. The loads tested were 100 through 1,000 N for a 4, 5, and 6 mm support with ribbing (Fig. 2). Our data shows the contact pressure across the gasket array with the smallest contact pressure occurring in the center of the device (left panel, contact pressure map of the gaskets) and highest on the edges with a minimum pressure of 0.06 MPa (4 mm fixture, Fig. 2A middle panel) with a maximum pressure of 18.6 Pa (6 mm fixture, Fig. 2A right panel). The pressure near both the edges and the center are higher with an increase in effective fixture thickness adjusted using a trapezoidal rib across the top of the fixture. Additionally, Fig. 2B displays these trends, but with respect to the resultant deformation of the gaskets and the fixture plate. The left panel shows an exaggerated bending due to the clamps on each end, although, the middle panel shows why the rib thickness must be optimized; as a 4 mm thickness separates the gasket away from the well (16 μm at 1000 N) in the center while a 5 mm and 6 mm thick rib enable clamping in the center (−8 μm and −32 μm respectively). The edge deformations are not significantly different near the loads (all approach −188 μm). Finally, the strain on the gaskets is mapped in Fig. 2C (left panel) and the center and edge strains for each load and thickness are plotted in the middle and right panels respectively.

**Figure 2.**
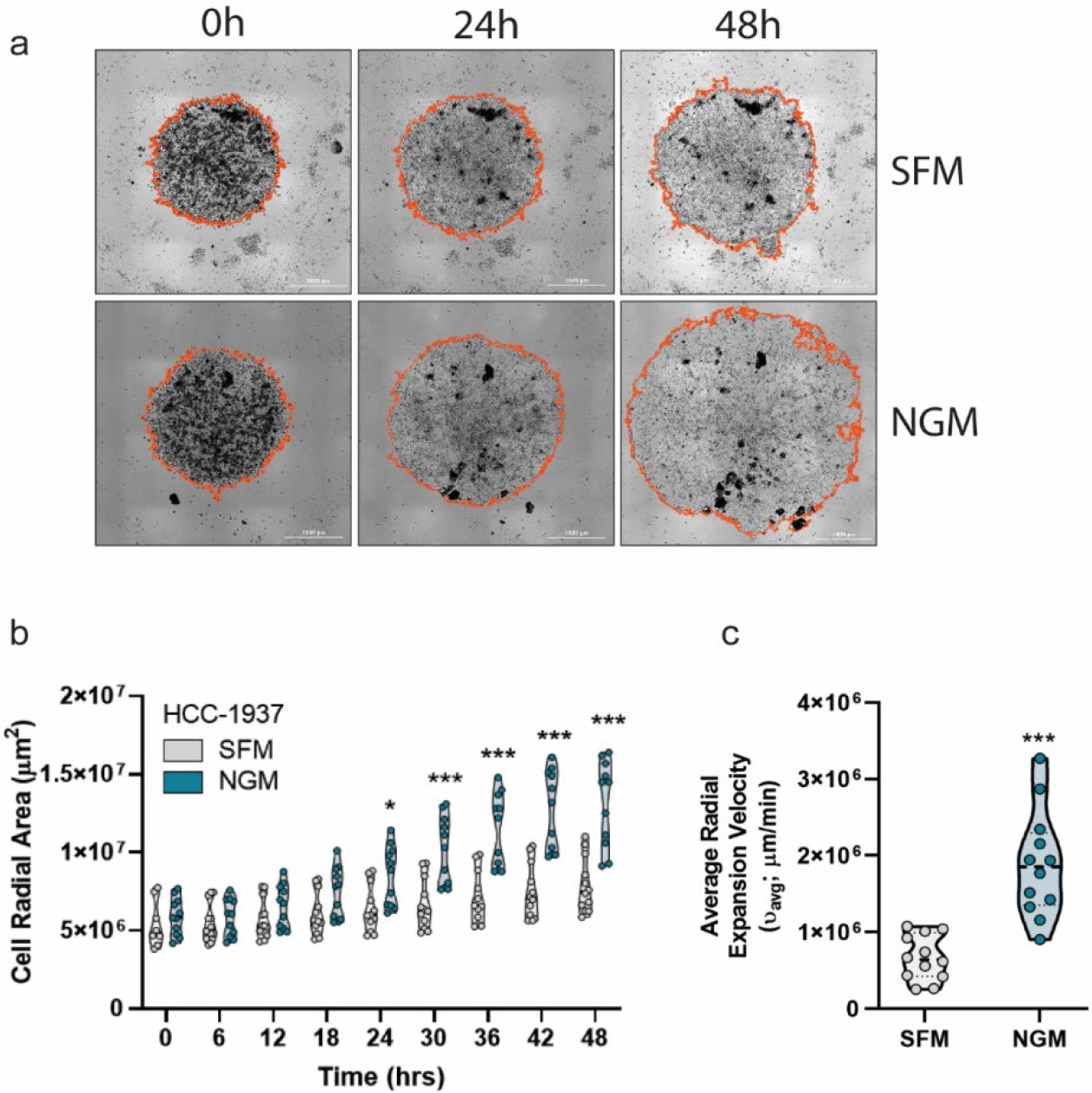
Breast cancer cell model HCC-1937 radial migratory responses and analysis. a) Representative images and Gen5 algorithm identification of HCC-1937 cells (orange ring) migrating in response to serum components in normal growth media (NGM) versus serum free media (SFM) conditions at the indicated time points. Scale bar; 1mm. b) Absolute cell radial area growth analysis of HCC-1937 cells in SFM vs NGM conditions. c) Radial expansion velocity measurements in SFM vs NGM conditions. Experiments displayed are representative of at least 2 independent biological replicates. Technical replicates in data shown is n≥4; statistical significance determined by two-way ANOVA. ***p<0.001

### Device Prototype Analysis

Based upon the results from FEA modeling (Fig. 2), the device was submitted for fabrication. The final platform/product as fabricated is shown in Fig. 3A. Gaskets clamped against a 24 well plate is shown in Fig. 3B. To validate the sealing and performance of the device, MDA-MB-231 breast cancer cells were seeded into the device as described in the “*Device Implementation and Experimental Approach*” section of this manuscript. Radial cell monolayer post seeding can be observed in Figure 3C. Upon deposition of cell media into each well of the device, we observed several instances of bubbles formation near the gasket-cell culture plate interface resulting in improper radial cell monolayer formation (Fig. 3D). To avoid the formation of bubbles at this interface, we find that application of vacuum (whole device placed inside desiccator bell jar; see *Device Implementation and Experimental Approach* subsection for further detail) greatly reduced bubble formation, significantly enhancing the reproducibility of the assay.

**Figure 3 –.**
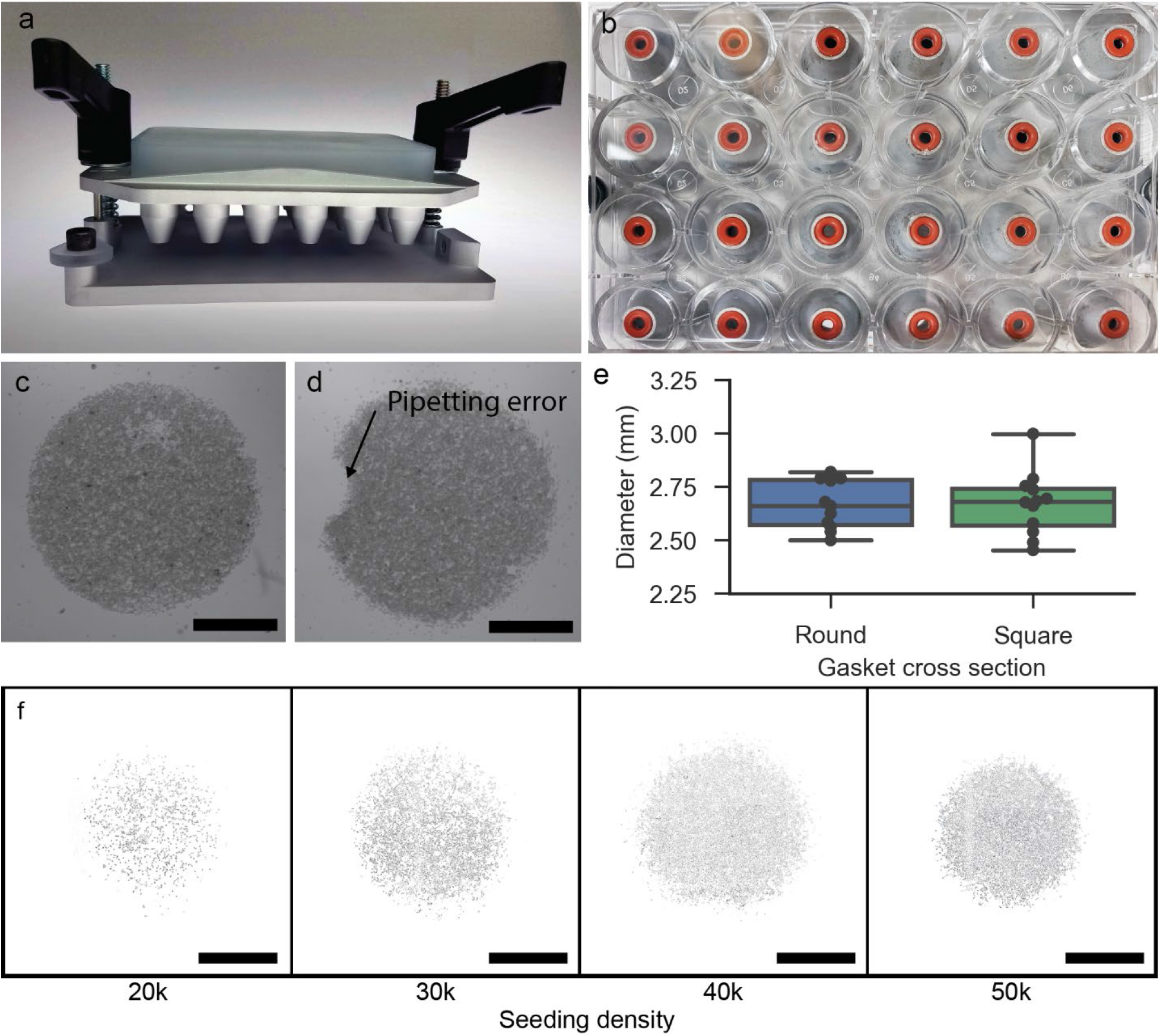
Validation and performance of the fabricated radial migration assay. a) Photograph of the assembled device as fabricated. b) Bottom up view of the gaskets sealed across the face of each well in a 24-well plate. c) Example of MDA-MB-231 breast cancer cells seeded onto the face of a well plate. Scale bar; 1 mm. d) Example of MDA-MB-231 breast cancer cells seeded onto the face of a well plate when a bubble was introduced into the edge of the device. Scale bar; 1 mm. e) Comparison of variability and differences in the diameter of cell monolayers as seeded into the device when either round or square cross section gaskets were used. f) MDA-IBC-3 cells seeded at differing densities. Scale bar; 1mm.

Next, we sought to determine the reproducibility of the size of the radial cell monolayer across the device. First, gaskets with a round cross sections were compared against gaskets with square cross sections. We suspected a square cross section may have lower variability between experiments their constant cross-sectional area when in contact with the cell culture plate. The resulting diameters of the cell spot were measured by brightfield microscopy (Fig. 3E). Variability is not significantly different between the two cross sections, and we chose the more readily available round cross section gaskets for all remaining experiments. The average diameter of radial cell monolayers was 2.66 mm with a standard deviation of 0.11 mm (n=24) (Fig. 3E). The maximum diameter was 2.81 mm, and the minimum diameter was 2.5 mm. Given the specified gasket diameter is 2.56 mm, this is consistent as the inner diameter of the rounded gasket will be slightly larger than the nominal gasket diameter (Fig. 3E blue). The gasket alignment plate is designed with a hard stop to ensure 20% of the gasket height is compressed.

### Proof of Principle Experimental Approach

To display the utility of our device and assay, we aimed to determine the migratory capacity of HCC-1937 human breast cancer cells in the presence or absence of growth serum. Using BioTek’s Gen5 analysis software, we utilized built in algorithms which provided robust identification and labeling of radial cell monolayers (Fig. 4A). We then quantified radial cell areas post expansion, measured every 6 hours. Expectedly, we observe increases in radial migration when HCC-1937 cells are provided growth serum as compared to serum free conditions (Fig. 4B). Kinetic analysis in the Gen5 platform permits for rate measurements as a function of radial expansion area measurements over time. Using this methodology, we can quantify radial expansion velocities (μm/minute) for each cell monolayer, averaged over the entire experimental analysis. Indeed, radial expansion velocities are a function of the area measurements but provide a more concise display of the data (Fig. 4C). Collectively, these data show the capacity for this assay to demonstrate significant changes in migratory rates dependent on the cellular environment. Indeed, this assay could be applied to observe phenotypic changes in response to drug regiments or monitor the role a genetic component may play in regulating migration, among various other scenarios.

## DISCUSSION

We have developed and presented a new device which adapts an important but underutilized migration assay, the radial migration assay. We believe the use of this method has been hampered by the limited tools available, the variability of the radial dimensions, significant technical difficulty in set up, and time required to perform extensive replicates. Here, we offer a high-throughput version of the radial migration assay, which is simplified, less laborious, and more user-friendly. Moreover, the high-accuracy, consistent placement of the cell spot enables imaging from a high-content, automated system.

The primary drawback of the system are the non-uniformity of the clamping pressure. Future design iterations will address this by modeling alterations to the clamping set up or perhaps modality. Consistently obtaining proper gasket contact pressure is currently challenging, although the implementation of a thread stop may aid in applying the correct clamping force on a more consistent basis. Additionally, while application of vacuum effectively removes bubbles formed by deposition of cell culture media, we do occasionally encounter bubbles forming at the gasket/culture plate interface. This technical error must be controlled for by the user by utilizing several technical replicates per experiment to allow for radial formation error (i.e. bubble formation and/or poor cell adherence). Lastly, significant care must be taken to utilize clean instruments (e.g. forceps) when placing or handling the gaskets such that dirt or oils are not deposited to the gasket face. Regardless of these prototyping issues, we believe in the merits of the approach as evidenced by our results, and technical error can be minimized by careful practices of the user.

Migration and motility are potentially the most crucial phenotypic characteristics cells intrinsically harbor. Cellular motility is an imperative aspect of tissue generation, organ formation, wound healing, inflammatory responses, among many other physiological functions that are essential for human survival (17–20). While many fields of biological research have significant interest in cellular migration, it is cemented as a hallmark of cancer and pinned as one of the key cellular functions which enable metastatic spread and ultimately patient mortality. Indeed, metastasis is the single most important identifier of patient mortality. Despite our recent advances in the development of targeted cancer therapeutics, patients who are diagnosed at late stage (i.e. III or later), have locally advanced tumors, or who have distant metastasis, have a very poor prognosis and low survial rates. This has highlighted the need for therapuetic regiments which are aimed at diminishing or managing metastasis via inhibition of cellular migration/invasion. Hence, we believe our device would be useful as an implementation to survey migratory and invasive phenotypes in response to metastates-targeted therapies. Additionally, our device and assay are useful in examining the role of molecular mediators of cellular migration/invasion post genetic modification of the target e.g. RNAi, overexpression, conditional expression, etc. Our application allows for a more high-throughput method to evalute migration/invasion in both 2D and 3D formats, collectively enhancing research progress at greater rates versus mutiformat migration assays.

As mentioned previously, an important differentiation from other 2D migration assays is such that the radial migration assay does not physically alter cells in the ways that a scratch or wound assay does. This mitigates responses from wounded or dying cells and offers information about the intrinsic, non-directional motility of a cell. Indeed, non-directional, inherently acquired cell motility by tumor cells is the precursor to metastatic invasion. Furthermore, there is room for adapting this system to understand how environmental cues or signals alter cellular motility. For instance, our system is designed to be utilized in a standard 24-well culture plate and could be paired with co-culture systems to understand how other immune cells of the tumor microenvironment (TME), e.g. tumor associated macrophages, impact cancer cell motility. Additionally, collagen or ECM matrix deposition on cells following removal of the device would allow for evaluation of cellular invasion. Future work may increase the number of cell types and other components that can be patterned using this method by fabricating custom gaskets inserts.

Collectively, we strongly feel our device can benefit a variety of researchers with wide-spanning interests related to cellular migration and motility. The ease of device use and implementation along with the simplicity of data acquisition and analysis permits a variety of users with diverse backgrounds to utilize our device and assay. While high-throughput imaging permits the acquisition of more robust kinetic data, users without this capability can simply obtain initial and endpoint images to determine migratory capacity. Our newly manufactured device coupled to a simple user interface permits the acquisition of migration data that was previously laborious and time consuming. Therefore, we strongly believe implementation of our device and assay can be of great benefit to any research application with vested interests in cellular migration or invasion.

## METHODS AND MATERIALS

### Device fabrication

The device was designed in Solidworks 3D design software. The files provided in the supplemental (S1-S6) were exported and uploaded to the digital fabricator Protolabs who then milled the parts out of 6061 aluminum with a tolerance of ± 0.005 inches. Three parts were fabricated by Protolabs Machining division including the gasket fixture plate which holds the gaskets in place (S1), the base for aligning the top plate to a well plate (S2), a lock to hold the well plate in place (S3). The parts were then tapped, edges broken and bead blasted for finishing. The final part was a lid designed to prevent contamination of the sample was fabricated by Protolabs 3D printing division out of the WaterShed XC 11122 material (S4). The aluminum parts were anodized by Alpha Metal Finishing in a hard coat grey anodization. The remainder of device components were purchased from McMaster-Carr. The components are provided in tabular form the supplementary information for those that which to use the device (S6). The purchased components include two clamps (McMaster, 6385K110), two springs (McMaster, 9434K820), two 1/8” diameter alignment pins (McMaster, 97395A454), one set screw (McMaster, 91658A154), two ¼-20 threaded rods 4 inches long (McMaster, 90322A652), one ¼-20 ½” long screw for locking the well plate in place (McMaster, 91251A527), 005 size silicone gaskets (McMaster, 1173N005), square cross section 005 size silicone gaskets (McMaster, 1182N005).

### Cell Models

Breast cancer cell models MDA-MB-231 and HCC1937 were acquired from ATCC and maintained in Gibco RPMI-1640, 10% FBS, 5μg/mL gentamycin, 2mM L-glutamine, and 1X antibiotic-antimycotic. Cells were cultured at 37°C and 5% CO_2_. MDA-IBC-3 cells were generously provided by Dr. Wendy Woodward (M.D. Anderson) and maintained in Gibco DMEM-F12, 10% FBS, 5μg/mL gentamycin, 1μg/mL insulin, 1μg/mL hydrocortisone, and 1X antibiotic-antimycotic. Cells were cultured at 37°C and 10% CO_2_.

### Radial Device Preparation and High Content Microscopy

Cells were seeded at 5.0×10^4^ per well of the radial migration device in a 24-well cell culture dish (Denville Scientific, NJ, USA). Cells were maintained at 5% CO_2_ and 37°C overnight (~16hrs) to permit adherence to 24-well culture dish. Radial device was removed, cells washed 3X with sterile PBS (Gibco) to remove non-adherent cells and/or debris and maintained in 500μL complete media. Images were taken on a BioTek BioSpa/Cytation5 automated incubator high-throughput imaging platform (BioTek, VT, USA) every 6 hours for 2 days. BioSpa incubator system and Cytation5 imager were maintained at 37°C and 5% CO_2_.

## CONTRIBUTIONS

ACL and CRO conceptualized the study. CRO designed the radial migration device. ACL, CRO, PP, and TW performed experiments. AL performed the high-throughput imaging and Gen5 analysis. CRO performed the FEA and modeling. ACL and CRO analyzed data. CRO and ACL wrote the manuscript. JAY and SDM provided critical input to the conceptual device and experimental design. SDM financially supported the research.

## ACKNOWLEDGEMENTS

Funding for this project is supported by the Breast Cancer Research Foundation and Rogel Cancer Center core grant NIH-P30-CA046592-29, P30CA046592 (SDM, AL, CRO), T32CA009676 (CRO), by the METAvivor Foundation (SDM) and the Breast Cancer Research Foundation (CRO, AL,SDM).

## Notes

#### Summary of Updates

We have recently updated some of the language associated in figure 1, added new data (Fig. 3F), and made figure 4 more concise and updated the axis to reflect the correct units of measurement. Additionally, we bolstered the discussion section and added important references in the text.

## REFERENCES

1. Lauffenburger DA, Horwitz AF. Cell Migration: A Physically Integrated Molecular Process 1996;84:359–369.

2. Valster A et al. Cell migration and invasion assays. Methods 2005;37(2):208–215.

3. Gilles C et al. Vimentin contributes to human mammary epithelial cell migration. [Internet]. J. Cell Sci. 1999;112 (Pt 2:4615–25.

4. McKenzie AJ, Campbell SL, Howe AK. Protein kinase a activity and anchoring are required for ovarian cancer cell migration and invasion. PLoS One 2011;6(10). doi:10.1371/journal.pone.0026552

5. Decaestecker C, Debeir O, Van Ham P, Kiss R. Can anti-migratory drugs be screened in vitro? A review of 2D and 3D assays for the quantitative analysis of cell migration. Med. Res. Rev. 2007;27(2):149–176.

6. Derda R et al. Multizone paper platform for 3D cell cultures. PLoS One 2011;6(5). doi:10.1371/journal.pone.0018940

7. Haridas P, McGovern JA, McElwain SDL, Simpson MJ. Quantitative comparison of the spreading and invasion of radial growth phase and metastatic melanoma cells in a three-dimensional human skin equivalent model. PeerJ 2017;2017(9). doi:10.7717/peerj.3754

8. Van Horssen R, Ten Hagen TLM. Crossing barriers: The new dimension of 2D cell migration assays. J. Cell. Physiol. 2011;226(1):288–290.

9. Todaro GJ, Lazar GK, Green H. The initiation of cell division in a contact-inhibited mammalian cell line. J. Cell. Physiol. 1965;66(3):325–33.

10. Liang C-C, Park AY, Guan J-L. In vitro scratch assay: a convenient and inexpensive method for analysis of cell migration in vitro. Nat. Protoc. 2007;2(2):329–333.

11. Shen Y et al. A General Approach for Fabricating Arc-Shaped Composite Nanowire Arrays by Pulsed Laser Deposition [Internet]. Adv. Funct. Mater. 2010;20(5):703–707.

12. Bianchi ME. DAMPs, PAMPs and alarmins: all we need to know about danger. J. Leukoc. Biol. 2007;

13. Krysko D V. et al. Immunogenic cell death and DAMPs in cancer therapy. Nat. Rev. Cancer 2012; doi:10.1038/nrc3380

14. Yarrow JC, Perlman ZE, Westwood NJ, Mitchison TJ. A high-throughput cell migration assay using scratch wound healing, a comparison of image-based readout methods. BMC Biotechnol. 2004;4:1–9.

15. Little AC et al. DUOX1 silencing in lung cancer promotes EMT, cancer stem cell characteristics and invasive properties. Oncogenesis 2016;5(10):1–11.

16. Little AC et al. IL-4/IL-13 stimulated macrophages enhance breast cancer invasion via rho-GTPase regulation of synergistic VEGF/CCL-18 signaling. Front. Oncol. 2019;9(MAY):1–13.

17. Yamada KM, Sixt M. Mechanisms of 3D cell migration [Internet]. Nat. Rev. Mol. Cell Biol. 2019;20(12):738–752.

18. Friedl P, Gilmour D. Collective cell migration in morphogenesis, regeneration and cancer. Nat. Rev. Mol. Cell Biol. 2009;10(7):445–457.

19. Fujimori T, Nakajima A, Shimada N, Sawai S. Tissue self-organization based on collective cell migration by contact activation of locomotion and chemotaxis. Proc. Natl. Acad. Sci. U. S. A. 2019;116(10):4291–4296.

20. Xiao Y, Riahi R, Torab P, Zhang DD, Wong PK. Collective Cell Migration in 3D Epithelial Wound Healing. ACS Nano 2019;13:1204–1212.

